# Large river eDNA sampling designs with remote-sensing-based clustering stratification

**DOI:** 10.64898/2026.04.29.720935

**Authors:** Shuo Zong, Robin Bauknecht, Hansjörg Seybold, Camille Albouy, Loïc Pellissier

**Author notes:** **Corresponding authors:** Robin Bauknecht, Shuo Zong. These authors contributed equally to this work (Shuo Zong, Robin Bauknecht).

## Abstract

Environmental DNA (eDNA) provides a powerful tool for biodiversity monitoring in large river ecosystems. However, current studies often rely on subjective site selection and lack systematic sampling designs. This can limit the ability to capture the full spectrum of environmental conditions that species depend on, thereby compromising sampling efficiency. To address this challenge, we propose utilizing remote sensing-based clustering for environmental stratification of sampling designs, thereby enhancing detection capabilities and increasing the objectivity of eDNA sampling. Using GBIF-based fish species distribution models and simulated distributions along the Danube, we demonstrate that this approach enhances detection efficiency compared to conventional random and regular sampling methods. To facilitate practical implementation, we developed a tool to help fieldwork planners of river sampling campaigns automatically apply this method and select stratified sampling sites without the need for extensive data processing. Finally, we demonstrate that eDNA detection occurred most frequently within the range of 0–20km downstream of the expected modeled distribution of species, suggesting that the diffusion of the signal should be further considered in the sampling design process. Our findings highlight the potential of remote sensing-based stratification to create more efficient and objective sampling designs but suggest that sampling design should be further combined with hydrological information to optimize cost-efficient sampling. The development of standard and robust sampling protocols will help advance more cost-effective eDNA-based biodiversity monitoring in riverine ecosystems.

## 1 Introduction

Rivers are essential components of the Earth’s natural landscape, supporting both ecological and human systems. They provide critical habitats for aquatic and terrestrial organisms (Lasso et al., 2016; Paetzold et al., 2005) and supply vital ecosystem services, such as freshwater and food, to human societies (Begossi et al., 2019; J. Stephen Lansing, 1998; Xu et al., 2023). The functionality of these systems is strongly dependent on a healthy aquatic species community (Febria et al., 2015; Hooper et al., 2005). However, the aquatic biodiversity of rivers is under severe threat from alterations such as the introduction of invasive species, dam construction, and land use modifications (Leuven et al., 2009; Vörösmarty et al., 2010; Wu et al., 2019). Understanding and monitoring the biodiversity of river ecosystems is essential for effective conservation and, thereby, for ensuring the sustained provision of the critical services that rivers offer to both nature and humans (Darwall et al., 2018; Miqueleiz et al., 2023). Sampling designs should be designed in a way that optimizes the cost-efficient detection of species and allows for the best attribution to the drivers of biodiversity within rivers.

Environmental DNA (eDNA) sampling has emerged as a tool to quickly survey and assess biodiversity in freshwater ecosystems, aiding in the improvement of monitoring schemes (Baird and Hajibabaei, 2012; Pawlowski et al., 2021). eDNA refers to a mixture of intracellular DNA from living cells and extracellular DNA from sources such as skin and hair, which is directly extracted from environmental samples such as water (Pawlowski et al., 2020). Combining high-throughput DNA sequencing with taxonomic reference libraries then allows for the rapid and comprehensive identification of the taxa from which the DNA originates (Deiner et al., 2017; Ruppert et al., 2019). This enables the rapid detection and identification of species without the need for direct observation or physical capture, thereby facilitating ecosystem assessment across large spatial scales (Cilleros et al., 2019; Deiner et al., 2016). Further, eDNA is recognized as a relatively cost-effective method with high detection capacity, including the ability to identify rare and elusive species (Boivin-Delisle et al., 2021; Jerde et al., 2011; Mächler et al., 2014; Polanco Fernández et al., 2021). Therefore, eDNA is progressively gaining prominence in assessing large-scale freshwater biodiversity (Altermatt et al., 2020; Lyet et al., 2021; Prié et al., 2023). However, while the laboratory procedure has matured to yield reliable detections, the sampling in eDNA studies remains poorly designed, and the selection of sampling locations rarely follows an optimized sampling design (Altermatt et al., 2023).

Minimizing effort and cost while optimizing the quantity and quality of data collected is a crucial aspect of a spatially detailed biodiversity assessment (Carraro et al., 2021). Despite significant research efforts to establish common standards for laboratory protocols (e.g., Coutant et al., 2021), there has been comparatively less focus on refining sampling procedures and data collection methods in the field (Altermatt et al., 2023). Dickie et al. (2018) found that 95 % of studies collected over three years used subjective eDNA sampling designs that did not provide critical methodological information. Conversely, only a few studies enhanced objectivity by randomly placing sampling sites or selecting them based on evident characteristics, such as the deepest point of the sampling area (Dickie et al., 2018). While these methodologies provide a general overview of species presence, they may struggle to capture the spatial variability of environmental factors that influence habitat-specific distributions of organisms within the sampling area (Zhang and Zhang, 2012). Consequently, this can lead to biased, inefficient, or incomplete assessments of the species communities under examination (Anderson, 2001). Since sampling errors often significantly exceed other analytical errors (Zhang and Zhang, 2012), improving sampling strategies is critical. One approach is stratified sampling, a statistical technique that enhances the representativeness of collected data by dividing the study area into subgroups, or strata, based on shared ecological characteristics. By ensuring that each subgroup is adequately represented, stratified sampling reduces bias and improves biodiversity assessments (Hirzel and Guisan, 2002; Lohr, 2021). Moreover, this systematic approach enhances objectivity and reproducibility, which are often lacking in eDNA sampling site selection (Dickie et al., 2018). However, its implementation requires detailed knowledge of on-site conditions, which may be difficult to obtain before fieldwork begins, adding complexity to the planning process.

Satellite remote sensing (hereafter, remote sensing) provides high-resolution, gap-free observations of the Earth’s surface. It can serve as a valuable tool for acquiring information about environmental conditions in the sampling area (Ozesmi and Bauer, 2002; Zong et al., 2024). A wide range of ecological properties, including vegetation indices, surface temperature, water quality parameters, and hydrological characteristics, can be acquired through remote sensing (Crippen, 1990; Malakar et al., 2018; Olmanson et al., 2008; Schmugge et al., 2002; Sims et al., 2006; Vanhellemont, 2020). By utilizing this data, researchers can create detailed maps of key environmental variables before fieldwork, facilitating more systematic planning of sampling efforts. In large river eDNA sampling, remote sensing can help classify water bodies into different environmental strata based on both aquatic and surrounding terrestrial characteristics. Ecological sampling studies have shown that methods such as random stratified sampling can allow the collection of datasets that can provide a stronger association with environmental parameters (Williams and Brown, 2019), which can, in turn, help generate more robust spatial models for, e.g., conservation applications. The use of ecological strata for rivers derived from remote sensing can inform the placement of sampling sites, ensuring a more systematic and representative selection. Despite its potential, this approach remains underutilized in eDNA research (Dickie et al., 2018), and its advantages over conventional methods, such as random sampling, require further evaluation, especially in the context of riverine systems.

eDNA has emerged as a powerful tool for biodiversity monitoring, yet its transport dynamics in large river systems present both challenges and opportunities for species detection (Pont et al., 2018). Pont et al. (2018) demonstrated that detection distance varied from a few km in a small stream to more than 100 km in a large river. In large river environments, eDNA can travel considerable distances downstream from the point of origin, leading to the detection of species beyond their occupied range (Brantschen et al., 2024). This downstream transport has significant implications for the design of eDNA sampling strategies in large rivers, as it can enhance species detection but also requires careful interpretation of results to avoid misattributing species distributions. By accounting for eDNA transport in sampling designs, researchers can optimize detection efforts, improving biodiversity assessments while minimizing false absences (Carraro et al., 2023). Understanding how eDNA moves through river networks is therefore essential for refining monitoring protocols and maximizing the efficiency of large-scale aquatic surveys (Pont et al., 2024).

In this study, we address this gap by developing a novel methodology that integrates remote sensing data with k-means clustering to create environmentally stratified river eDNA sampling designs. Our approach aims to enhance the efficiency and representativeness of eDNA surveys by capturing the environmental heterogeneity of river systems (Ward and Tockner, 2001). We achieve this by clustering river segments based on key environmental variables derived from remote sensing, ensuring that sampled sites encompass the full range of habitat conditions. Unlike traditional strata definitions, clustering provides a more objective and data-driven method for identifying distinct environmental zones. To evaluate its effectiveness, we apply our approach to the Danube River, Europe’s second-longest river, renowned for its rich biodiversity and ecological significance (Liška et al., 2021; Sommerwerk et al., 2022). Using data from the Global Biodiversity Information Facility (GBIF) and remote sensing-derived environmental variables, we develop species distribution models (SDMs) for fish species inhabiting the Danube and generate simulated fish distributions to provide a controlled evaluation framework. We then compare our stratified sampling designs against random and systematic sampling using both the SDMs and simulated distributions to assess improvements in detection efficiency. By integrating remote sensing data and advanced clustering techniques, our study aims to enhance eDNA-based biodiversity monitoring in river ecosystems.

## 2 Methods

### 2.1 Remote sensing-based sampling stratification

We employed k-means clustering to create environmentally stratified sampling designs. This unsupervised machine learning algorithm partitions data into *k* clusters by iteratively assigning points to the nearest centroid and updating centroids until convergence (Jain et al., 1999; Lloyd, 1982). Applied to key environmental variables from remote sensing, it allows objective classification of the river into distinct strata, ensuring representative sampling by selecting an equal number of sites from each cluster (Maggini et al., 2002). Given the limited knowledge about variations in the eDNA concentration across a river transect (Macher and Leese, 2017), we solely optimize the sampling site positioning along a river and consequently simplify rivers as lines. To facilitate the practical implementation of clustering-based stratified sampling, we developed a GEE-based tool that allows researchers to select a river segment of interest and obtain river strata, as well as optimized sampling sites without complex data processing. The river polylines in the tool are retrieved from the HydroSHEDS Free Flowing Rivers data set available on GEE (Grill et al., 2019; Lehner et al., 2008). Users can customize the methodology to suit their specific needs by selecting environmental variables that they suspect influence the distribution of the target taxa in their study area. Additionally, users can adjust parameters such as the number of clusters, the number of sampling sites, and the minimum distance between sampling sites. The tool outputs the coordinates of the optimized sampling sites and provides visualization options to display the clusters and selected sites on a map. We generated fish SDMs (Guisan and Zimmermann, 2000; Guisan et al., 2017) as the first approach to evaluate the efficiency of the approach. SDMs provide a controlled environment that allows for repeated testing, ensuring a thorough and reliable evaluation. We chose the Danube River as our case study due to its well-documented species records and the abundance of fish observations available throughout its course (Schiemer et al., 2004). We considered all 102 species described in the extensive fish species list of the Danube described by Schiemer et al. (2004).

#### 2.1.1 Species occurrence data

We compiled occurrence data for these species from GBIF (GBIF.Org User, 2024). We initially extracted all observations of the species in the 10 countries the Danube traverses and then filtered this data to retain only occurrences within a 1500 m buffer zone on each side of the main river channel polyline. This buffer was chosen to encompass the maximum width of the Danube, approximately 1500 m (International Commission for the Protection of the Danube River, 2024), as well as the surrounding areas since many GBIF observations are located in the confluence zones of smaller rivers flowing into the Danube. Although not directly on the Danube, these observations are still applicable as they reflect the species’ presence in nearby, interconnected habitats, enhancing our predictions for the main channel. To address the lack of sites with confirmed species absences inherent in the presence-only data from GBIF, we employed pseudoabsence sampling (Guisan et al., 2017). We created a probability surface along the water pixels of the Danube using geographic inverse distance weighting (IDW) through the `geoIDW` function from the `dismo` package (Hijmans et al., 2011). IDW is a spatial interpolation method that estimates unknown values based on their proximity to known values (Bartier and Keller, 1996), here assigning higher weights to locations closer to known occurrences. Specifically, we used all species observations as well as an equal number of spatially random absences to create the interpolation surface (Descombes et al., 2022). This approach mitigates geographical sampling bias and ensures higher pseudoabsence sampling probabilities near observed occurrences, where intensive inter-specific sampling effort increases confidence in true species absences. From this probability surface, we sampled an equal number of absences and presences for each species, maintaining ecological relevance by limiting pseudoabsence placement to water areas.

#### 2.1.2 Environmental predictors

Several variables characterizing fish habitat suitability in rivers were calculated using remote sensing imagery. The river water surface temperature (RST), a critical determinant of fish metabolic rates (Gillooly et al., 2001), was computed using thermal infrared sensor data from Landsat-8 (Vanhellemont, 2020; Zong et al., 2024). The Secchi Depth (SD), a proxy for water transparency, was calculated using products from the Multispectral Instrument (MSI) aboard Sentinel-2 and the Landsat-8 Operational Land Imager (OLI) (Drusch et al., 2012; Page et al., 2019). Fish habitat suitability is not only influenced by aquatic but also by terrestrial factors, as the riparian vegetation influences the availability of terrestrial insects and fruits, provides essential shade, and creates diverse microhabitats for fish to live in (Beltrão et al., 2009; Cetra and Petrere JR., 2007; Montag et al., 2019; Vannote et al., 1980). Therefore, we calculated the Enhanced Vegetation Index (EVI) and forest canopy height (CH) which provide information about vegetation structure (Lang et al., 2023; Sims et al., 2006). Further, gross primary productivity (GPP), a proxy for ecosystem productivity, and total evapotranspiration (EP), quantifying the release of water vapor from the forest canopy and surroundings, were computed from MODIS products (Gebremichael and Barros, 2006; Kumagai et al., 2005). The slope of the water surface along the river channel was extracted as it governs the flow dynamics of the river and thus potentially impacts fish migration patterns and spawning behaviors (Farr et al., 2007; Mercado-Silva et al., 2012). The elevation of the site was extracted as rivers can exhibit fish species assemblage changes along the upstream-downstream gradient (Carvajal-Quintero et al., 2015; Oberdorff et al., 1993). The analysis also incorporated the Human Modification Index (HMI) which quantitatively measures human impact on the surrounding environment, encompassing various anthropogenic alterations (Kennedy et al., 2019). We considered satellite imagery covering the five-year period 2019–2023 to assess the long-term habitat condition whilst avoiding picking up short-term fluctuations. Predictor values were extracted for all presence and absence sites, using the mean within a 5 km circular buffer zone (Hijmans, 2024; Pebesma, 2018; Pebesma and Bivand, 2023; R Core Team, 2013). This buffer size captures broader ecological influences on mobile freshwater fish and has proven effective for modeling fish species distributions along the Rhone River, which is geographically close to the Danube (Zong et al., 2024).

**Table 1:** Overview of the remote sensing variables used in the analysis. Computations were performed in the Google Earth Engine cloud computing platform.

| Abbreviation | Description | Resolution |
| --- | --- | --- |
| RST | Mean river water surface temperature (Vanhellemont, 2020) |  |
| SD | Mean Secchi depth within the buffer (Page et al., 2019) |  |
| EVI | Mean enhanced vegetation index (Sims et al., 2006) |  |
| GPP | Mean gross primary productivity |  |
| CH | Mean canopy height (Lang et al., 2023) |  |
| EP | Mean total evapotranspiration |  |
| Slope | Mean river surface slope (Farr et al., 2007) |  |
| Elev | Mean elevation |  |
| HMI | Mean human modification index (Kennedy et al., 2019) |  |

#### 2.1.3 Fish SDM fitting and evaluation

Given the limited sample size, we chose to employ GLM models (Dobson and Barnett, 2018), using a binomial link to handle presence/absence data. Such models can predict the probability of a species being present at a site by modeling the relationship between the predictors and the log odds of presence. To avoid multicollinearity, predictors with high correlations (Pearson correlation *>* 0.7) were removed across all species (Dormann et al., 2013). The `step` function (R Core Team, 2013) was used to iteratively select the best set of predictors for each species, ensuring an optimal model fit with minimal complexity.

To evaluate the performance of our models, we performed 5-fold cross-validation for each species (Kohavi, 1995). The folds were created using the `createFolds` function from the `caret` package, ensuring each fold had a roughly equal proportion of presences and pseudoabsences (Kuhn and Max, 2008). For each species, models were iteratively trained on four-folds, including stepwise model selection (`step` function; R Core Team, 2013), and used to make predictions on the remaining fold. We calculated the average Cohen’s Kappa (Cohen, 1960) and True Skill Statistic (TSS) (Allouche et al., 2006), two commonly used evaluation metrics, across all folds to assess the predictive model’s performance. We used the `optimal.thresholds` function of the `PresenceAbsence` package (Freeman and Moisen, 2008a) to determine the optimal threshold for classifying predicted probabilities into presence or absence, based on maximizing the Kappa statistic (Freeman and Moisen, 2008b). Models with a performance score below 0.2 for either Kappa or TSS were excluded from further analysis, as such models would only show slight agreement with the data (Allouche et al., 2006; Munoz and Bangdiwala, 1997), and we wanted to ensure reliability in testing our sampling design methodology on the SDMs. Additionally, species with only one or two occurrences (resulting in only 2 or 4 sites when combined with pseudoabsences) were excluded because our cross-validation procedure is not applicable for such limited data. Using the models selected through cross-validation, we predicted species distributions along the main river channel polyline.

### 2.2 Simulated fish distributions

To complement our empirical SDMs, we simulated theoretical fish distributions based on the environmental conditions along the Danube. Specifically, we generated species-specific logistic response curves using the five most influential predictors identified in the SDMs. A total of 54 response curves were created to match the final number of SDMs considered. Each simulated species was categorized as rare, intermediate, or standard, with intercept values randomly assigned based on this classification. Rare species were assigned lower intercepts, resulting in lower overall occurrence probabilities, while common species were assigned higher intercepts, leading to greater predicted prevalence. For each species, we then assigned random slope values corresponding to the five environmental predictors, representing their response to each of those factors. The linear predictor was computed as the weighted sum of the scaled ecological variables and then transformed using the logistic function to yield occurrence probabilities ranging from 0 to 1, providing a theoretical distribution of species presence across the study area.

### 2.3 Sampling design efficiency evaluation

To assess the performance of our stratified sampling approach in comparison to random and regular sampling, we applied these methods to both the empirical SDMs and simulated species distributions. Stratified sampling was based on the five most influential predictors from the SDMs, under the assumption that these predictors, on average, have the most substantial influence on the distributions. Using these variables, we partitioned the river into ten clusters, each representing similar environmental conditions. We then selected an equal number of sampling sites from each cluster to ensure balanced coverage. For regular sampling, we created evenly spaced sampling locations along the river. To introduce variation across replicates, we shifted the sampling points by a distance equal to one percent of the segment length for each replicate. Random sampling was performed by selecting sites thoroughly at random along the river. Each sampling approach was evaluated across a range of sample sizes, from 10 to 100, in increments of 10, with 100 replicates generated for each sample size to ensure robust evaluation. We used two different detection approaches to determine species presence at each site. The first, a probabilistic detection approach, models the probability of detection as the product of the predicted species probability and a detection rate of 0.1. The second approach, a threshold-based method, classifies species presence using species-specific thresholds set at 95 % of the maximum predicted probability. These conservative choices for detection probability and thresholds aimed to highlight differences in the methods, as a higher detection rate or lower threshold would lead to more uniformly high detection rates across all sampling sizes, thereby diminishing the differences. However, while adjusting these values influences the absolute number of detected species, the overall patterns remain consistent.

### 2.4 Transport of the eDNA signal

To examine the alignment of GBIF-based Species Distribution Models (SDMs) with eDNA water sample data, we utilized a published eDNA dataset from the Danube River, which includes 24 sampling locations with two eDNA sample replicates per location. Those 24 sites were sampled on the Danube River close to the average hydrological conditions, with sufficient distance between the sites to avoid the potential influence of eDNA transported downstream from one site to the next. Each water sample was collected with a peristaltic pump inside a disposable sterile tube and was directly filtered on the boat through a cross-flow filtration capsule (VigiDNA 0.45 µm, SPYGEN), and its volume was measured. At the end of each filtration, the water in the capsule was drained, and the capsule was refilled with conservation buffer CL1 (SPYGEN) to prevent eDNA degradation. The capsule was then kept at room temperature until DNA extraction. The eDNA laboratory workflow (extraction, amplification using “teleo” primers, high-throughput sequencing) was performed following a previously described protocol (Pont et al., 2018). We employed OBITOOLS to assign raw sequences to the species level. Initially, we extracted common fish species from both the environmental DNA data and GBIF data. For each common species range from the SDM, we identified the nearest downstream eDNA sampling locations containing signals of these species. We recorded the shortest distance between the eDNA signal and the corresponding species range. Given the complex morphology of the river due to its meandering nature, we aimed to achieve accurate distance measurements along the river channel. To do this, we converted the entire river into milestone points at 100-meter intervals. We then assigned these milestones to both eDNA sampling locations and SDM ranges, allowing us to calculate the distances between them easily.

## 3 Results

### 3.1 SDM evaluation

Our analysis identified 54 fish species for which distributions along the Danube River can be reliably modeled. These models, built on a total of 5321 occurrences along the river (average of 98 per species), demonstrated a mean Cohen’s Kappa of 0.607 *±* 0.192 and a TSS of 0.612 *±* 0.191 in cross-validation, indicating good predictive performance (Allouche et al., 2006; Munoz and Bangdiwala, 1997). Initially, 67 of the 102 species known to occur in the Danube had observations along the river. However, six species were excluded due to insufficient sample sizes for cross-validation, and seven species were removed due to evaluation scores below 0.2. This process resulted in the final set of 54 species that provided reliable SDMs for further analysis.

### 3.2 Sampling design efficiency

Stratified sampling covered a broader range of environmental variance along the river compared to both random and regular sampling (Figure 1). This pattern was consistent for both small (*n* = 10) and large (*n* = 100) sample sizes. While random sampling captured more environmental variation than regular sampling, the latter exhibited the lowest variance across replicates (Figure 1).

**Figure 1:**
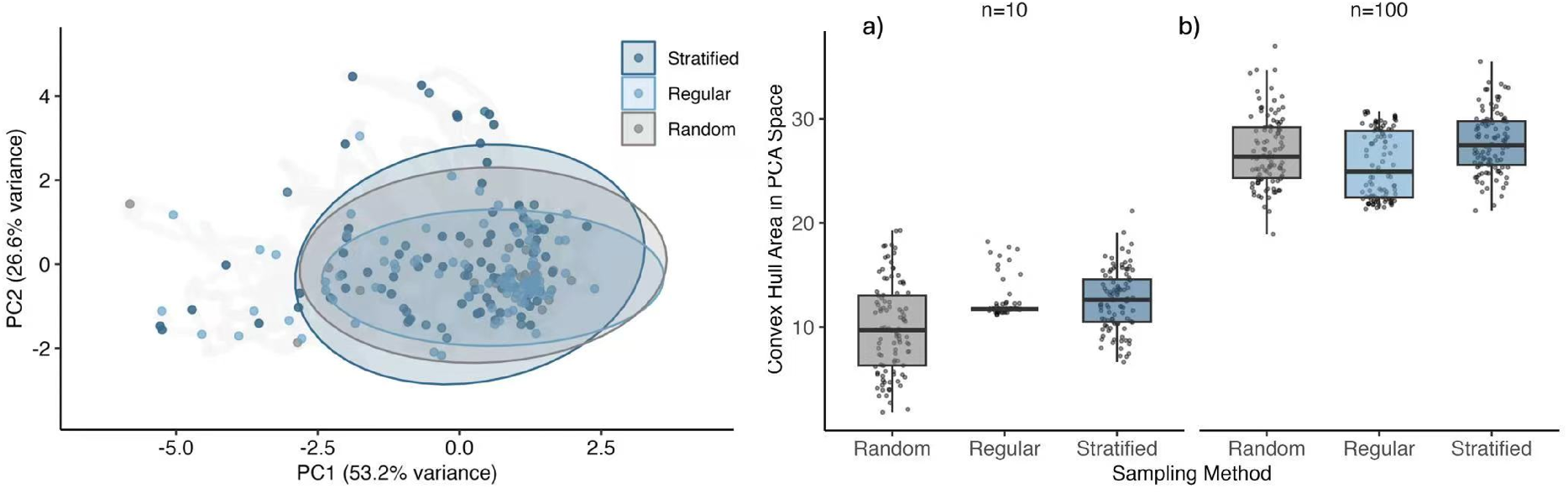
Principal components of sampling locations by different sampling designs. Left Panel: PCA ordination plot showing the distribution of environmental conditions across all river pixels (gray points) in relation to 100 sampling points selected using random, regular, and stratified sampling. The PCA was performed on the five most influential predictors in the SDMs. Variables were standardized to ensure equal contribution of each variable to the PCA. The ellipses indicate the 95 % confidence intervals for each sampling method, illustrating the environmental space captured by each approach. Right Panel: Boxplots of the convex hull areas in PCA space, calculated using the first two principal components (PC1 and PC2), for each sampling method across 100 independent runs and two different sample sizes (*n* = 10 and *n* = 100). The convex hull area represents the environmental coverage achieved by each sampling strategy, with larger areas indicating broader environmental representativeness. The results highlight the differences in coverage between stratified, regular, and random sampling approaches.

The stratified sampling design also detected more unique species than random and regular sampling across most tested sample sizes and detection approaches (Figure 2). This advantage was particularly pronounced under the threshold-based detection method. On average, compared to random sampling, for a given sample size, environmentally stratified sampling identified 0.47 *±* 0.42 more species with probabilistic detection and 3.23 *±* 0.19 more species with threshold detection. Compared to regular sampling, for a given sample size, environmentally stratified sampling identified 0.79 *±* 0.44 more species with probabilistic detection and 1.47 *±* 1.30 more species with threshold detection. Across all methods, increasing the sample size led to the detection of more species. Notably, regular sampling had the lowest variance across replicate runs, particularly under the threshold-based approach (Figure 2).

**Figure 2:**
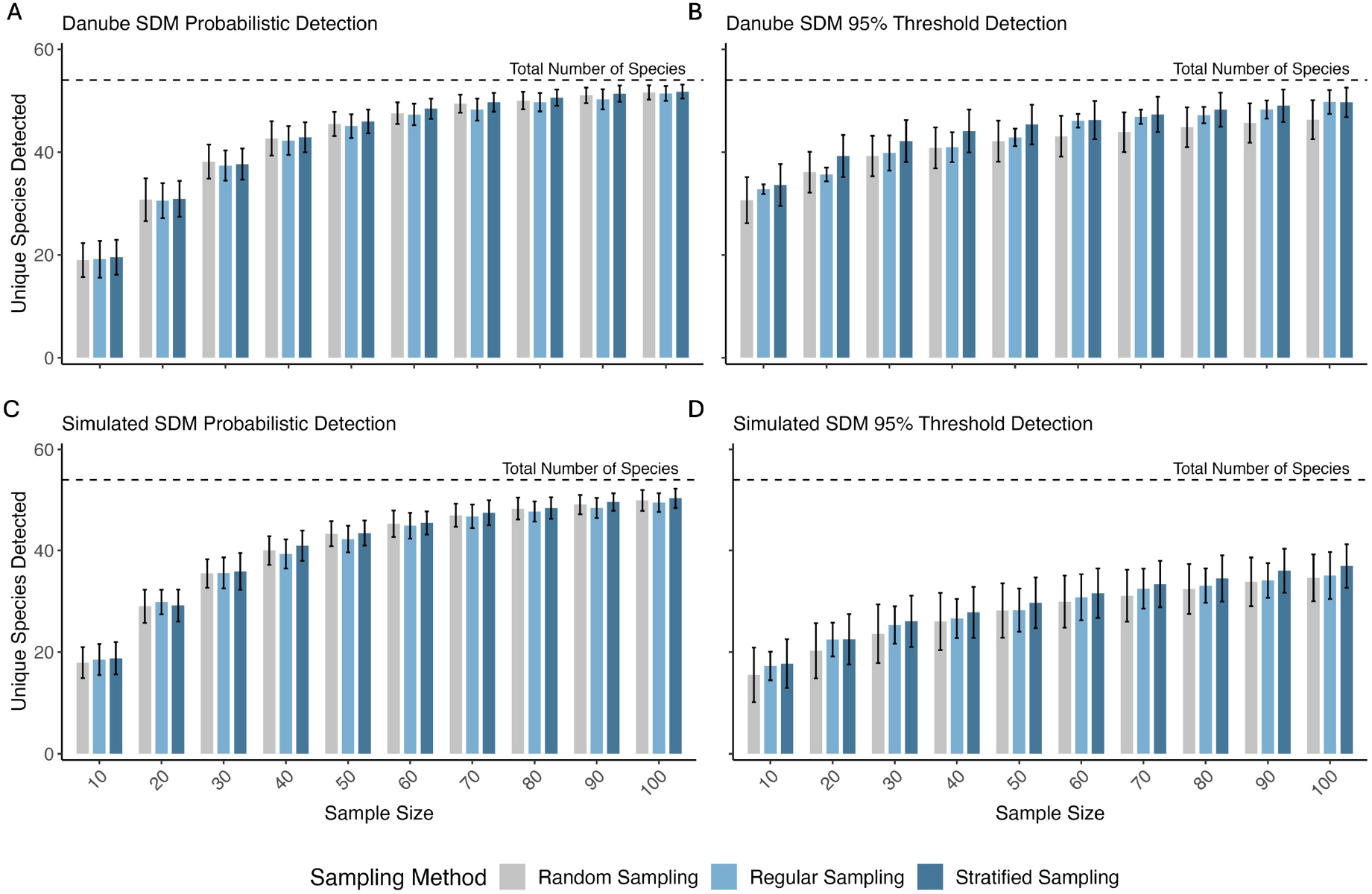
Comparison of the number of unique species detected using random, regular, and stratified sampling designs at various sample sizes. Each bar represents the mean number of unique species detected across 100 replicate runs, with error bars indicating the standard deviation of species counts among replicates, reflecting detection variability. The dashed line denotes the total number of species in the study area (54 species). Panels **A** and **B** show results from testing on GBIF-based fish SDMs along the Danube River using probabilistic and 95% threshold detection, respectively. Panels **C** and **D** present results from testing on simulated species distributions under the same detection criteria.

Stratified sampling did not increase the mean species presence probability at any sample size (Figure 3). However, it did enhance the maximum species presence probability across low (*n* = 10), intermediate (*n* = 50), and high (*n* = 100) sample sizes (Figure 3). This increase in the maximal predicted presence probability across samples was most pronounced at small sample sizes and decreased as the sample sizes increased.

**Figure 3:**
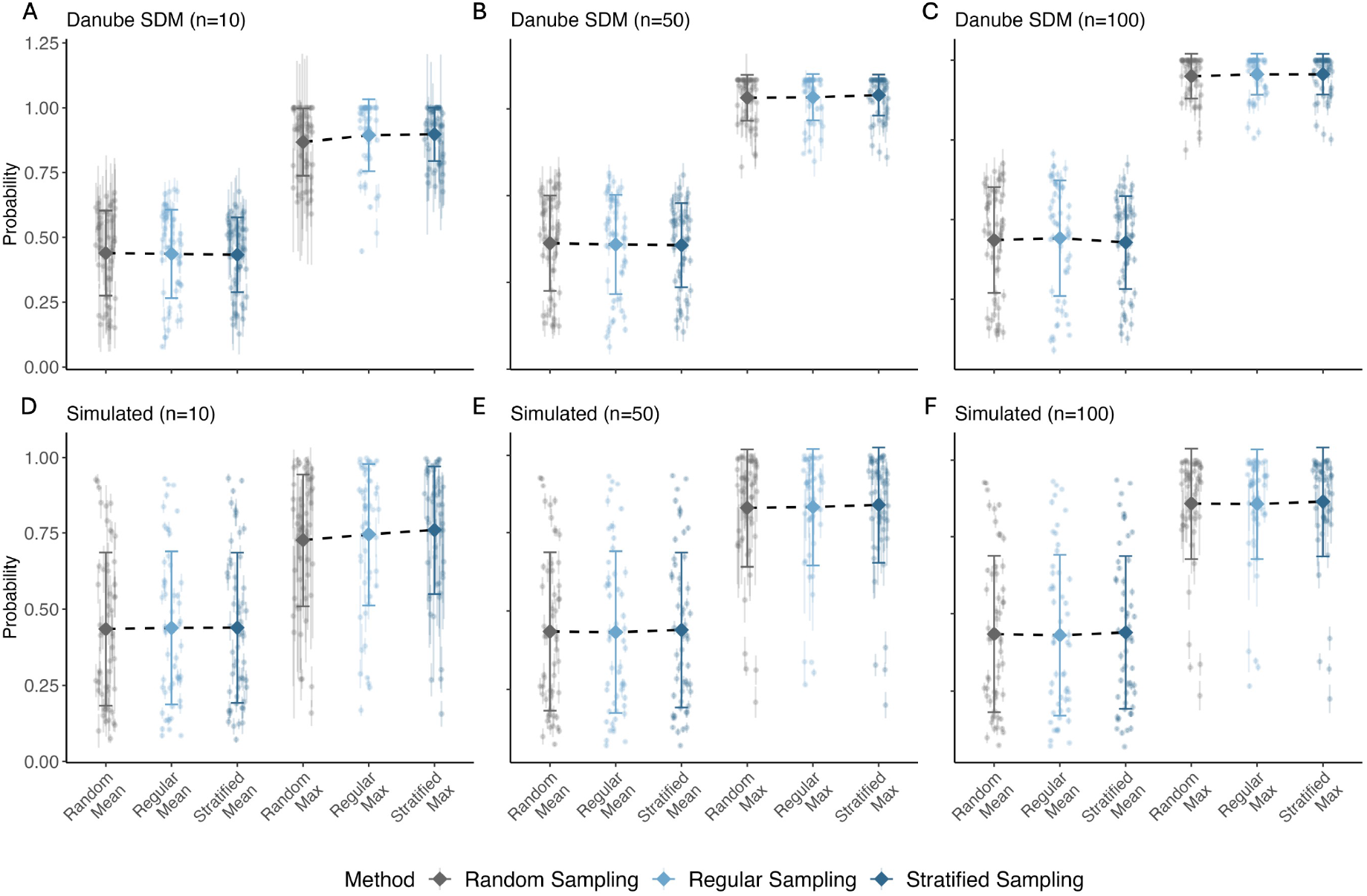
Overview of mean and maximal species presence probabilities for the Danube SDMs. (**A–C**) and simulated distributions (**D–F**) across different sample sizes: *n* = 10 (**A/D**), *n* = 50 (**B/E**), and *n* = 100 (**C/F**). Each point represents a single species, with error bars indicating the standard deviation across 100 replicates per sample size. Larger points and error bars represent population-level averages and deviations. “Mean” and “Max” refer to the average and maximum presence probability of a species across all sampled sites within a given run.

**Figure 4:**
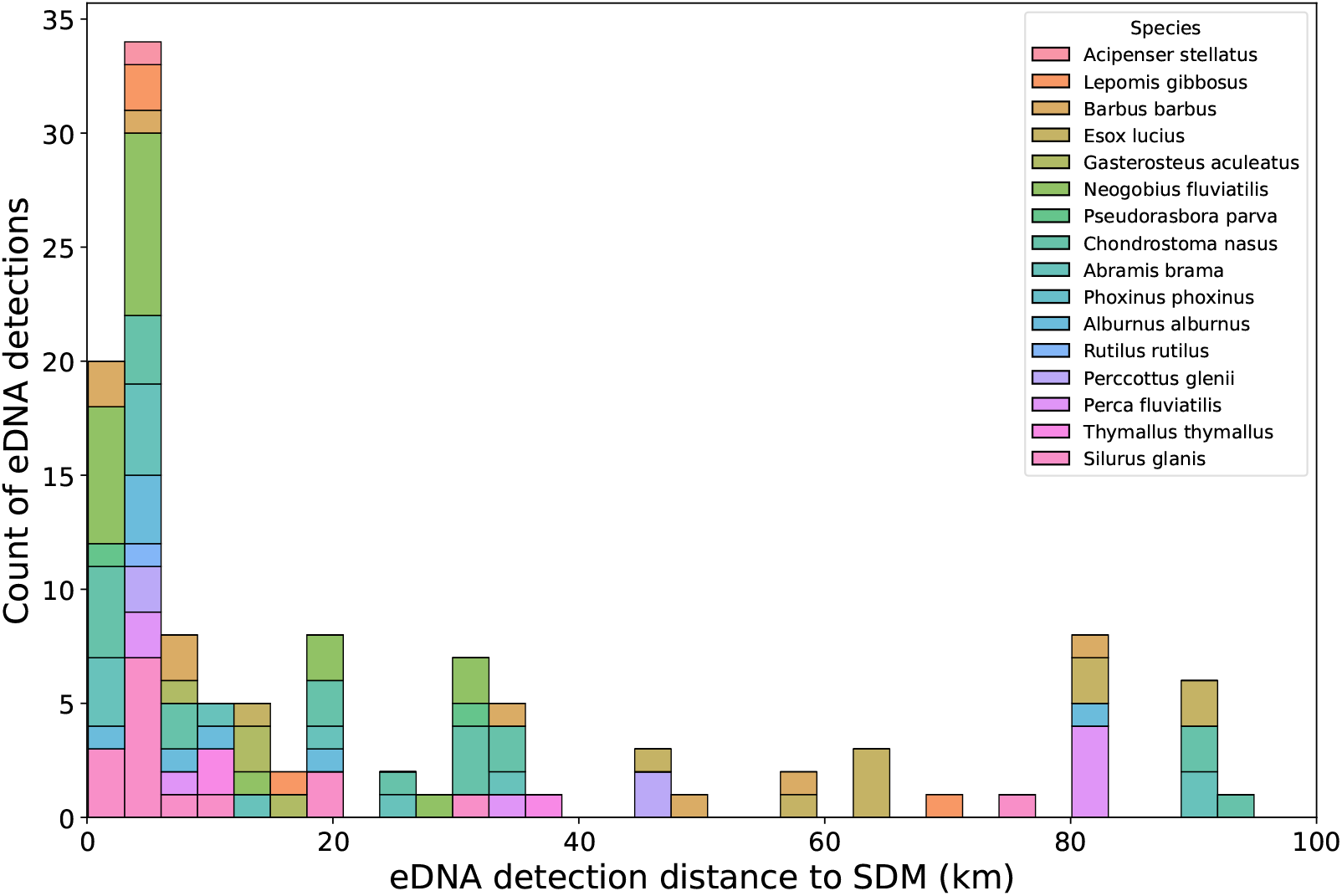
Comparison of species signals detected by eDNA samples in the Danube and GBIF-based fish species range from SDMs. Values on the x-axis indicate that the eDNA signal was found downstream from the fish species range. Count on the y-axis indicates how many eDNA samples detected the species at a given range of distance downstream. The color represents different species.

### 3.3 eDNA signal transport observation

By comparing the observed species detection from eDNA water samples with the predicted suitable range for each species, we identified areas where SDM predictions align with eDNA detections and areas where discrepancies occur. We observed the transportation of eDNA signals over a range of distances from 0 to 100 km downstream, with most of the distances occurring within the range of 0 to 20 km. We found that 56.23 percent of the distances were comprised between 0 and 5 km, indicating that generally, the eDNA detected is geographically close to its potential source. This observation is crucial for understanding the dispersal patterns of eDNA and its implications for species distribution modeling (SDM). We also observed that the distances were significantly different among the species considered (*p <* 0.05).

## 4 Discussion

We evaluate a random stratified approach to improve eDNA sampling in river systems by leveraging remote sensing-based environmental classification. We aimed to provide a practical tool that practitioners can use to improve their sampling designs for eDNA studies, thereby facilitating more effective and objective biodiversity monitoring and attribution to environmental drivers of change. Within the river system, a random stratified sampling approach offers two key advantages over traditional methods: first, it ensures better coverage of environmental conditions along the river (Figure 1), which in turn improves species detection rates (Figure 2); second, it introduces more objective, data-driven sampling designs into a field that has often relied on subjective decisions (Dickie et al., 2018). These advancements are particularly crucial in the context of river ecosystems, where environmental variability is high, and subjective or random site selection for sampling may overlook significant habitat differences, leading to incomplete or biased biodiversity assessments. By combining remote sensing data with clustering techniques, our method addresses these challenges, resulting in a more systematic, reproducible, and effective approach to eDNA sampling.

Stratified sampling outperformed random and regular sampling on both SDMs and simulated distributions for both detection approaches, demonstrating its effectiveness (Section 3.2, Figure 2 A–D). The increase in unique species detected across small, intermediate, and large sample sizes highlights its advantages for both resource-limited monitoring efforts—restricted to a few locations along the river—and extensive sampling campaigns. This enhanced detection capacity likely stems from the broader environmental coverage provided by stratified sampling across different sample sizes (Figure 1 C). By capturing a greater range of ecological variance along the river, stratified sampling ensures that more distinct habitats are included, increasing the likelihood of detecting species with specific niche requirements (Hirzel and Guisan, 2002). Supporting this explanation, we found that stratification increases the maximal observed species probability rather than the mean. This occurs because sites selected through stratification are more likely to include optimal habitat conditions for certain species. However, the mean probability remains unaffected, as some sites are more suitable for certain species. In particular, this effect is strongest at small sample sizes, where random selection is less likely to capture diverse habitat types. Regular sampling generally outperformed random sampling and exhibited the lowest variance across replicate runs of all methods. This stability likely results from its structured site selection, which avoids the random effects inherent in both fully random sampling and the stratified method’s within-stratum selection. Beyond efficiency, stratification provides a systematic framework that can enhance objectivity in a field that heavily relies on subjective sampling site choices. Using clustering to perform stratification has the advantage that the definition of strata does not depend on subjective thresholds, ensuring that sampling designs are grounded in objective, data-driven insights rather than subjective criteria (Jain et al., 1999).

The Danube fish SDMs we developed for evaluating the sampling approaches, on average, showed good performance as reflected by their Cohen’s Kappa and TSS scores (see Section 3.1). We were able to properly model 54 of the total 102 species known to historically inhabit the Danube (Schiemer et al., 2004), which is a positive outcome given that only 72 fish species were detected in the extensive Joint Danube Survey 4 (JDS4) conducted across 13 countries in 2019. The discrepancy between historical records and recent surveys likely reflects ongoing habitat loss and fragmentation due to rapid human modifications along the Danube (Kováč, 2015). Among the environmental variables, elevation emerged as the most important predictor across the models (Figure 5). This upstream-downstream gradient in fish biodiversity, indicated by our finding that lower elevations are particularly favorable for fish species, is supported by research from Balon (1964), who observed the highest species diversity in the Lower Danube, attributed to the influence of migratory species from the Black Sea. Furthermore, the Upper Danube is more heavily impacted by human activities, such as dam construction and increased navigation, which have been shown to have a negative effect on fish populations (Kováč, 2015; Schiemer et al., 2004). The importance and response direction of elevation in our models aligns with these ecological observations, lending further credibility to the developed models. To further strengthen our assessment, we introduced simulated species distributions, providing an independent, controlled framework to test the efficiency of different sampling strategies. These simulated species were designed to represent a range of ecological responses by assigning species-specific relationships to key environmental predictors. The consistency of results between SDMs and simulated distributions (Figure 2) suggests that the benefits of stratified sampling apply to both real-world species and theoretical species distributions, thereby reducing the risk that findings are dependent on the particular set of species modeled.

**Figure 5:**
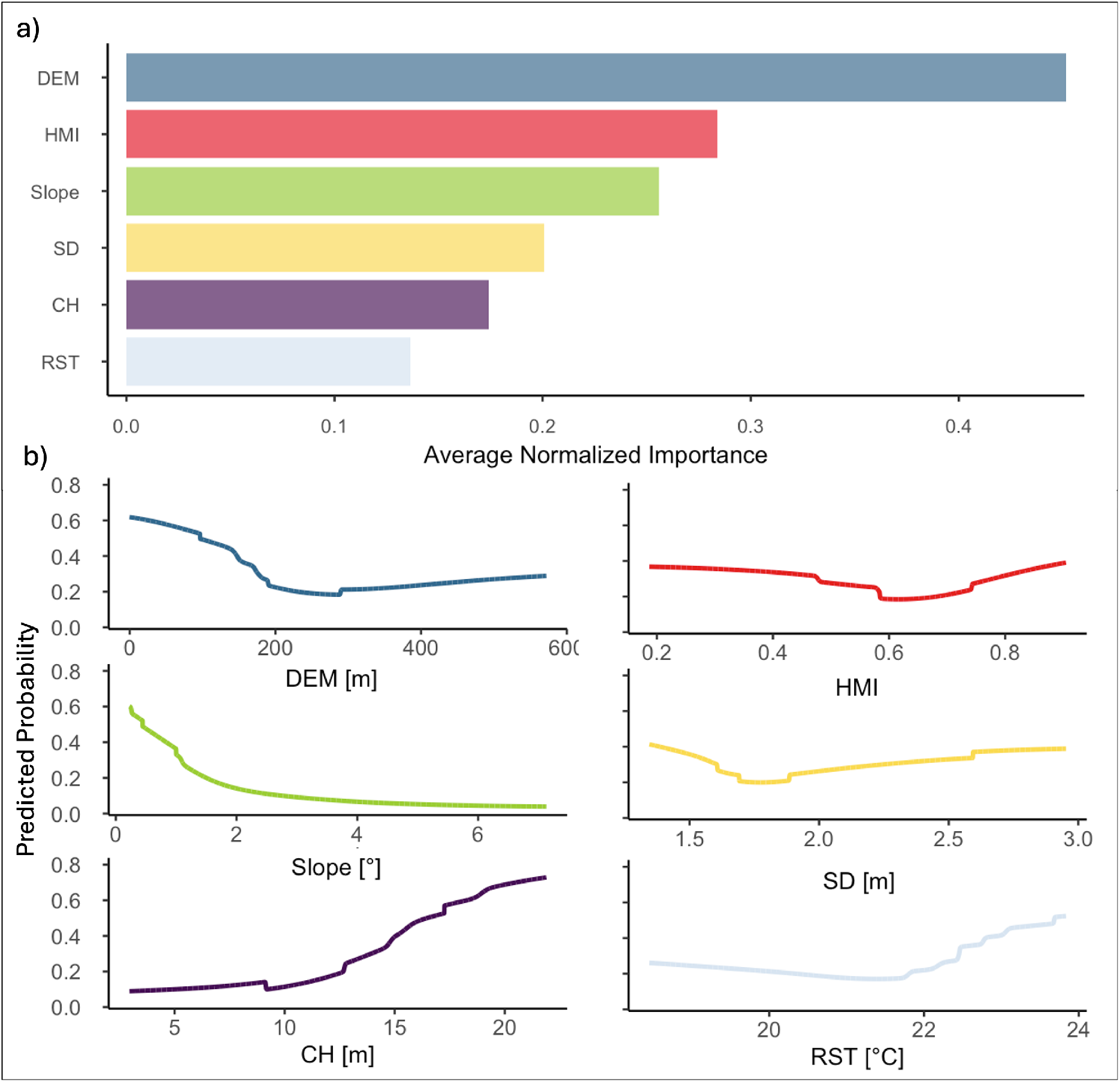
Average normalized variable importance across the 54 selected SDMs (a) and corresponding accumulated response curves (b).

While our methodology was tested on fish species within the Danube River, the approach is broadly applicable across various river systems and taxa (e.g., mammals). However, the success of this approach depends on making informed, context-specific decisions regarding the selection of environmental variables, the number of clusters, and the overall sampling strategy. Expert knowledge of the target river system remains crucial for tailoring these parameters effectively. For instance, in highly diverse river systems that traverse multiple biomes, a greater number of clusters may be necessary to capture distinct habitat types. Thus, while the methodology provides a robust framework, its optimal implementation requires careful consideration and adjustment based on the unique characteristics of each river system.

Environmental DNA (eDNA) transport dynamics in large rivers significantly influence biodiversity detection, posing substantial methodological and interpretative challenges (Deiner et al., 2016; Pont et al., 2018). Due to the downstream transport and dilution effects characteristic of large fluvial systems, eDNA signals can become spatially displaced from their sources, potentially leading to misinterpretations of species distribution and abundance and the calibration of inaccurate biodiversity models (Carraro et al., 2020a; Jane et al., 2015). In our analyses, we detected frequent downstream transport from 0 to 5 km, and possibly up to 100 km. Consequently, organisms detected at a sampling site may reflect upstream populations rather than local biodiversity, complicating ecological assessments and conservation efforts (Shogren et al., 2017). To mitigate these challenges, sampling designs should strategically incorporate longitudinal and lateral sampling points, paired with temporal replicates, to differentiate local versus transported eDNA signals (Laporte et al., 2020; Seymour et al., 2018). Furthermore, integrating hydrological modeling to predict eDNA transport and decay rates may greatly enhance the accuracy of species detection and spatial interpretation, ultimately informing management decisions more effectively (Carraro et al., 2020b; Harrison et al., 2019). Our analyses detected an effect of species on eDNA transport, and possibly, species with particular biology might shed more DNA that gets transported further away. Together, the sampling design should carefully consider both the theoretical framework and the empirical realities of eDNA transport dynamics and potential species-specific variations to ensure reliable biodiversity assessments and informed ecological interpretations.

Although our approach shows promising potential for improving eDNA sampling, several key limitations should be acknowledged. The methodology was tested only on the Danube River, while providing valuable insights and does not capture the full diversity of river systems worldwide (Penn, 2001). Rivers may vary in their environmental gradients, species assemblages, and hydrological dynamics, meaning that the method’s effectiveness may differ across contexts. However, the adaptability of our approach, particularly in terms of selecting environmental variables and the number of clusters, allows it to be tailored to other river systems, provided that expert knowledge is applied. The use of SDMs in our evaluation introduces another limitation. These models are built on presence-only data, which can be biased by uneven sampling efforts and gaps in species records (Beck et al., 2014). Additionally, our testing was based on cross-validation within the same dataset, which, while rigorous, does not replace the value of independent validation (Berrar, 2019). Unfortunately, an independent testing set was not available, and field testing would be ideal. eDNA sampling captures only a small fraction of the water column, which itself is subject to variability and bias (Mathieu et al., 2020). Thus, while our evaluation might not fully eliminate these biases, it provides a realistic and controlled assessment of the method’s relative performance. Another challenge lies in our approach’s reliance on binary presence/absence maps without explicitly modeling detection efficiency. While modeling detection efficiency on the SDM could offer additional insights, it is a complex task fraught with high uncertainty, as it involves numerous variables that are difficult to predict especially when working with large river systems. Our approach partially mitigates this by using high detection thresholds and multiple runs. Although the absolute estimates of species detection might thus vary from real-world sampling, the relative advantages of stratified sampling should remain consistent, ensuring that our conclusions are valid.

## 5 Conclusion

Our research demonstrates the potential of environmentally stratifying river eDNA sampling designs. By ensuring comprehensive coverage of environmental gradients, stratified sampling enhances the detection effectiveness of eDNA surveys. Further, clustering-based environmental stratification is independent of subjective threshold setting and enables more systematic and reproducible site selections. By providing this tool, we aim to encourage the broader adoption of systematic, data-driven sampling methods in river eDNA studies, ensuring more effective use of limited resources for biodiversity conservation.

